# Novel mechanisms of efflux-mediated levofloxacin resistance and reduced amikacin susceptibility in *Stenotrophomonas maltophilia*

**DOI:** 10.1101/2020.07.18.210146

**Authors:** Punyawee Dulyayangkul, Karina Calvopiña, Kate J. Heesom, Matthew B. Avison

## Abstract

Fluoroquinolone resistance in *Stenotrophomonas maltophilia* is multi-factorial, but the most significant factor is production of efflux pumps, particularly SmeDEF. Here we report that mutations in the glycosyl transferase gene *smlt0622* in *S. maltophilia* K279a mutant K M6 cause constitutive activation of SmeDEF production, leading to elevated levofloxacin MIC. Selection of a levofloxacin-resistant K M6 derivative, K M6 LEV^R^, allowed identification of a novel two-component regulatory system, Smlt2645/6 (renamed as SmaRS). The sensor kinase Smlt2646 (SmaS) is activated by mutation in K M6 LEV^R^ causing over-production of two novel ABC transporters and the known aminoglycoside efflux pump SmeYZ. Over-production of one ABC transporter, Smlt1651-4 (renamed as SmaCDEF) causes levofloxacin resistance in K M6 LEV^R^. Over-production of the other ABC transporter, Smlt2642/3 (renamed SmaAB) and SmeYZ both contribute to the elevated amikacin MIC against K M6 LEV^R^. Accordingly, we have identified two novel ABC transporters associated with antimicrobial drug resistance in *S. maltophilia*, and two novel regulatory systems whose mutation causes resistance to levofloxacin, clinically important as a promising drug for monotherapy against this highly resistant pathogen.

## Introduction

Levofloxacin is one of only six antimicrobials where breakpoints have been defined by CLSI for use against the opportunistic pathogen *Stenotrophomonas maltophilia* (1) The drug of choice is trimethoprim-sulphamethoxazole, but there have been several trials and meta analyses pointing towards the promising potential of levofloxacin monotherapy (2-4).

Fluoroquinolone resistance (e.g. to ciprofloxacin, moxifloxacin, levofloxacin) in Gram-negative bacteria involves multiple mechanisms (5). In Enterobacteriaceae, mutations in the fluoroquinolone targets, the so-called quinolone resistance determining regions (QRDRs) of DNA gyrase and topoisomerase enzymes are prevalent in fluoroquinolone resistant isolates. But in non-fermenting bacteria such as *Pseudomonas aeruginosa*, mutations increasing the production of fluoroquinolone efflux pumps are more common (5). For *S. maltophilia*, QRDR mutations have never been seen in clinical isolates or laboratory selected fluoroquinolone resistant mutants (6). Production of Qnr proteins, which protect DNA gyrase from fluoroquinolones, is important for intrinsic fluoroquinolone MICs against *S. maltophilia*, e.g. the chromosomally-encoded SmQnr (7,8) whose production is controlled at the transcriptional level by SmqnrR (9,10). We have recently shown that loss of TonB in *S. maltophilia* elevates fluoroquinolone MIC, suggesting that drug uptake is at least partly TonB dependent (11) but the most abundant fluoroquinolone resistance mechanisms in *S. maltophilia* are efflux pumps. These include the ABC transporter SmrA (Smlt1471) (12) the MFS type transporter MfsA (13) and the RND pumps SmeJK (14) and SmeGH (15).

The most clinically important fluoroquinolone efflux pumps in *S. maltophilia* are the RND systems SmeDEF and SmeVWX. SmeDEF was first identified as being hyper-produced in isolates resistant to a range of antimicrobials (16). Hyper-production was shown to be due to loss-of-function mutation in the transcriptional repressor gene *smeT*, encoded immediately upstream of *smeDEF* (17). Interestingly, triclosan is a substrate for SmeDEF and binds SmeT, meaning that SmeDEF production is induced in the presence of this biocide (18). It has been suggested that internal signal molecules may exist in *S. maltophilia*, which also bind SmeT and control *smeDEF* transcription (19). The role of SmeVWX over-production in fluroquinolone resistance in *S. maltophilia* clinical isolates is also well documented, particularly in the context of levofloxacin resistance, and particularly in combination with other mechanisms of resistance (20-22).

The work presented here reports the identification of novel regulatory elements, including a novel two-component regulatory system, and a novel ABC transporter contributing to levofloxacin resistance in *S. maltophilia* and demonstrates the associations between increased levofloxacin and amikacin MIC, identifying the amikacin transporters responsible.

## Results

### Disruption of glycosyl transferase gene smlt0622 causes over-production of SmeYZ and SmeDEF efflux pumps, leading to elevated amikacin and levofloxacin MICs against S. maltophilia K279a

We have previously defined *S. maltophilia* acquired ‘resistance profile 1’ in mutants with reduced susceptibility to fluoroquinolones and tetracyclines (19). Two such mutants are K M6 and K M7, derived from the clinical isolate K279a by selection for reduced susceptibility to moxifloxacin (19). The MIC of ciprofloxacin was previously found by Etest to have risen from 2 µg.mL^-1^ against K279a to be >32 µg.mL^-1^ against K M7 and 12 µg.mL^-1^ against K M6 (19). According to semi-quantitative RT-PCR, both mutants over-express *smeDEF*, which encodes the efflux pump associated with resistance profile 1 (19).

Both K M6 and K M7 were recovered from storage and confirmed by disc testing to have reduced susceptibility, but not to the point of resistance, to minocycline and trimethoprim/sulphamethoxazole, according to CLSI breakpoints (1) (**Table 1**). The most clinically relevant change came for levofloxacin, where K M6 was found to have acquired intermediate resistance and K M7 was found to be resistant, based on MIC testing (**Table 2**).

**Table 1.** Susceptibility testing of *S. maltophilia* K279a and mutants selected for reduced fluoroquinolone susceptibility. Shaded values represent reduced zone diameters (≥5 mm relative to K279a). For Disc susceptibility, values reported are the means of three repetitions rounded to the nearest integer for the diameter of the growth inhibition zone across each antimicrobial disc (mm). Susceptibility (S) is defined using breakpoints set by the CLSI (1). Where no designation is given, there is no defined breakpoint. Abbreviations: CAZ, ceftazidime; MH, minocycline; CN, gentamicin; C, chloramphenicol; SXT, sulphamethoxazole/trimethoprim.

|  | Zone Diameter (mm) Across Disc<br>(µg in disc) |  |  |  |  |
| --- | --- | --- | --- | --- | --- |
|  | CAZ<br>(30) | MH<br>(30) | CN<br>(30) | C<br>(30) | SXT<br>(25) |
| **K279a** | 32 | 32 (S) | 22 | 25 | 27 (S) |
| **K M6** | 30 | 27 (S) | 23 | 23 | 22 (S) |
| **K M7** | 31 | 27 (S) | 21 | 22 | 22 (S) |
| **K M6 LEV<sup>R</sup>** | 30 | 27 (S) | 16 | 22 | 22 (S) |

**Table 2.** MICs (μg.mL^-1^) against *S. maltophilia* K279a and mutant derivatives. The CLSI susceptible and resistance breakpoints (1) for levofloxacin are ≤2 and ≥8 µg.mL^-1^. There are no breakpoints for amikacin. Values are modes of three repetitions.

|  | Levofloxacin MIC | Amikacin MIC |
| --- | --- | --- |
| K279a | 2 | 8 |
| K M7 | 8 | 8 |
| K M6 | 4 | 16 |
| K M6 *smeE* | $\leq 0.25$ | 16 |
| K *smlt0622* | 8 | 64 |
| K LEV 5 | 8 | >256 |
| K LEV 5 *smlt2646* * | 2 | 8 |
| K279a *ampR*<sup>FS</sup>/pBBR1MCS-4 | 2 | 16 |
| K279a *ampR* <sup>FS</sup> /pBBR1MCS-4:: *smlt2644-6* * | 4 | >256 |
| K LEV 5 *smlt2643* | 8 | 64 |
| K LEV 5 *smlt1651* | 4 | >256 |
| K LEV 5 *smeE* | 0.5 | >256 |

Whole envelope proteomics analysis confirmed previously reported (19) over-expression of *smeDEF* in these two mutants. There was a 1.5-fold upregulation of SmeDEF in K M6, and a 3-fold upregulation of SmeDEF in K M7 relative to the parental strain, K279a (**Figure 1A**). The statistically significantly increased amount of SmeDEF produced in K M7 versus K M6 explains why MICs of ciprofloxacin (19) and levofloxacin (**Table 2**) are higher against K M7 than against K M6. Indeed, disruption of *smeE* in K M6, K M7 or K279a reduced the MIC of levofloxacin to 0.25 µg.mL^-1^, confirming the importance of SmeDEF for levofloxacin non-susceptibility in both mutants.

**Figure 1.**
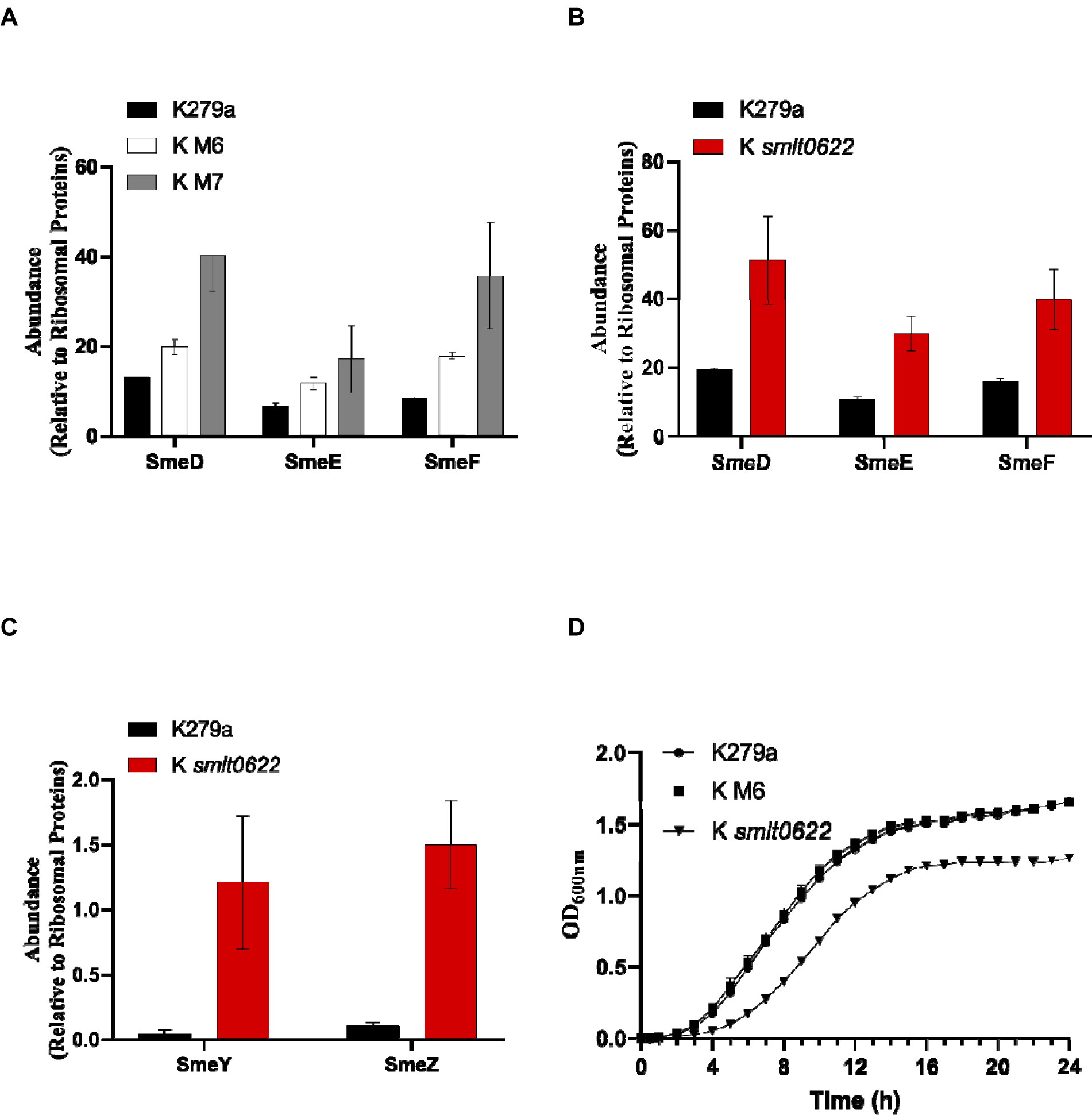
Role of glycosyl transferase Smlt0622 in controlling SmeDEF and SmeYZ efflux pump production. Protein abundance was measured using LC-MS/MS and normalised to the abundance of ribosomal proteins in cell extracts obtained from bacteria grown in NB. Data are mean ± standard error of the mean, *n*=3. Protein abundance in all mutants is statistically significantly different from the parent strain according to t-test (*p*<0.05). (**A**) SmeDEF production in the *smeT* loss-of-function mutant K M7 and the *smlt0622* point mutant K M6 versus the parent strain K279a (**B**) SmeDEF production in the *smlt0622* insertionally inactivated mutant versus K279a control. (**C**) SmeYZ production in the *smlt0622* insertionally inactivated mutant versus K279a control. (**D**) growth curve, in NB, of K279a, the *smlt0622* point mutant K M6 and the *smlt0622* insertionally inactivated mutant; growth based on OD_600_ was measured and presented as mean ± standard error of the mean.

K M7 has a loss-of-function mutation in *smeT*, but the mutation responsible for *smeDEF* over-expression in K M6 has not been defined (19). Whole genome sequencing revealed only one mutation in K M6, a single missense mutation relative to K279a, predicted to cause a Gly368Ala change in a putative glycosyl transferase encoded by the *smlt0622* gene. Glycosyl transferases are responsible for the addition of saccharides onto other biomolecules. Therefore, they can utilize various substrates and participate in myriad cellular functions. For example, cellular detoxification (23). Currently, there is no information about the specific role of the glycosyl transferase encoded by *smlt0622*.

To test whether the mutation in *smlt0622* is responsible for SmeDEF over-production in K M6, we insertionally inactivated *smlt0622* in its parent strain, K279a. Levofloxacin MIC was actually higher against K279a *smlt0622* than against K M6 (**Table 2**) and proteomics confirmed that SmeDEF production was higher in K279a *smlt0622* than in K279a, and higher even than in K M6, mirroring levofloxacin MIC (**Figure 1A, 1B, Table 2**). This led us to conclude that the Gly368Ala point mutant Smlt0622 enzyme in K M6 retains some activity. It is possible that Smlt0622 modifies a ligand that is the signal for SmeT de-repression or generates a ligand essential for SmeT repressive activity. Therefore, when the activity of Smlt0622 is reduced, the balance of ligand concentration is towards SmeT de-repression and *smeDEF* over-expression (**Table 2**).

We also noticed that the MIC of amikacin against K279a *smlt0622* was higher than against K279a (**Table 2**). This was explained by our observation from proteomics data that levels of SmeYZ, a known aminoglycoside efflux pump (24) were higher in K279a *smlt0622* than in K279a (**Figure 1C**). This was unexpected, because of previous data showing that SmeDEF over-production leads to reduced SmeYZ production (25); in this case K279a *smlt0622* over-produces both efflux pumps (**Figure 1**). One explanation is that the *smlt0622* mutation has a general effect on cellular physiology and that this stimulates SmeYZ production despite SmeDEF over-production. In support of this, we noted that K279a *smlt0622* grew slowly compared with K279a and the *smlt0622* point mutant K M6 (**Figure 1D**). We have recently reported that ribosome damage stimulates SmeYZ production in *S. maltophilia* (26) and so we hypothesise that slow growth activates a similar control system to ribosomal damage, stimulating SmeYZ production.

### ABC transporters controlled by the Smlt2645/6 two-component regulatory system contribute to levofloxacin resistance and elevated amikacin MIC

We next attempted to learn more about mechanisms of levofloxacin resistance in *S. maltophilia* by selecting a levofloxacin resistant mutant derivative of K M6. The resulting mutant, K M6 LEV^R^, presented a generally similar resistance profile to K M6 (**Table 1**) but had acquired levofloxacin resistance, as confirmed by MIC testing (**Table 2**). Interestingly, the mutant also had reduced susceptibility to the aminoglycosides gentamicin (**Table 1**) and amikacin (**Table 2**). Whole envelope proteomic analysis (**Table 3**) revealed upregulation of a bipartite ABC transporter (Smlt2642/3) in K M6 LEV^R^ versus K M6 (**Figure 2A**). We also noticed in the proteomics data that a putative two-component regulatory system (Smlt2645/6), encoded immediately adjacent to *smlt2642*/*3* on the chromosome, is also over-produced in K M6 LEV^R^ relative to K M6 (**Figure 2A**). According to whole genome sequencing, K M6 LEV^R^ has only one mutation relative to K M6, predicted to cause an Ala198Thr change in the over-produced sensor kinase Smlt2646. This putative Smlt2645/6 two-component system is therefore a good candidate for local activation of *smlt2642/3* ABC transporter operon transcription in K M6 LEV^R^.

**Table 3:**
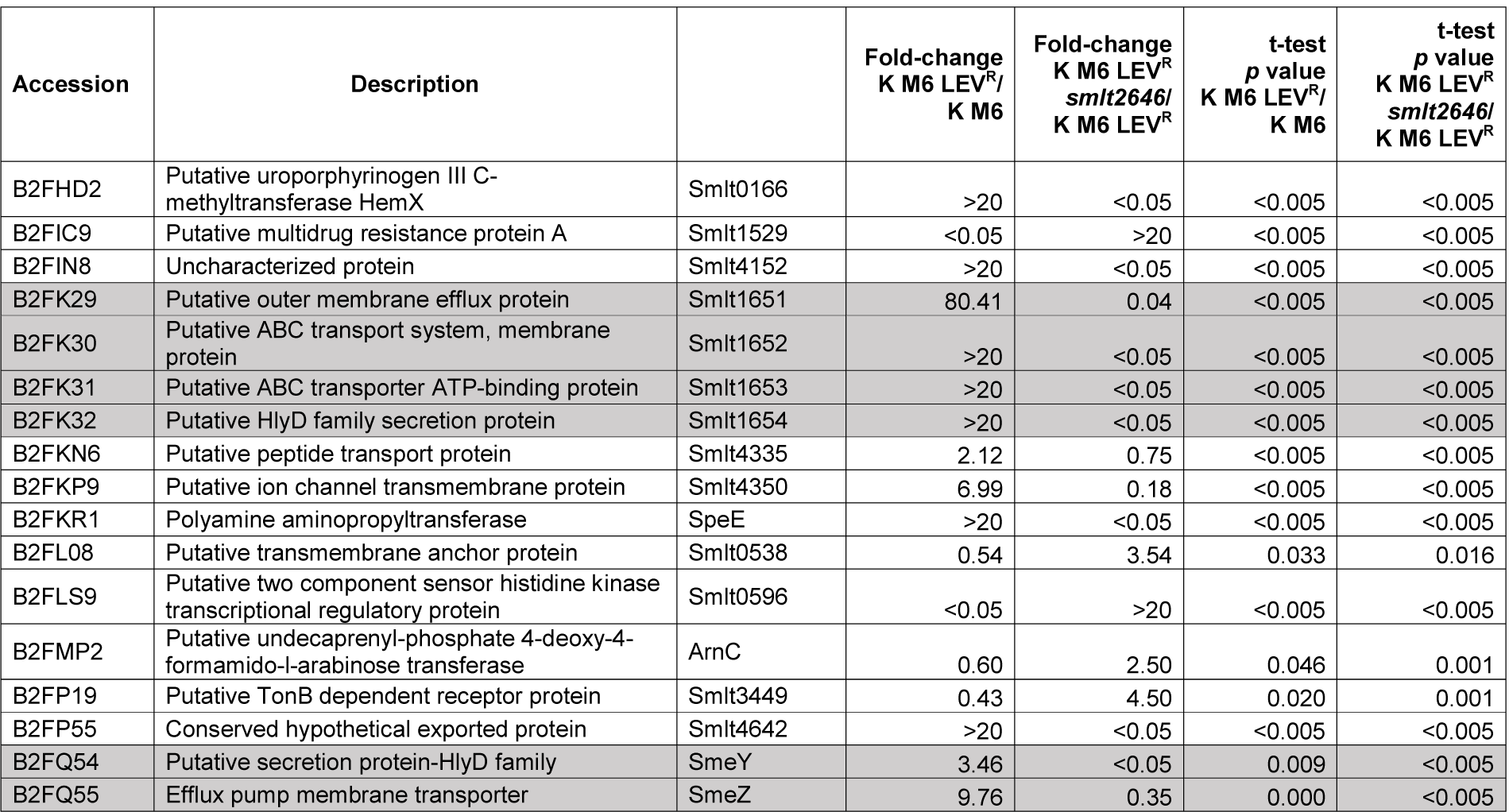
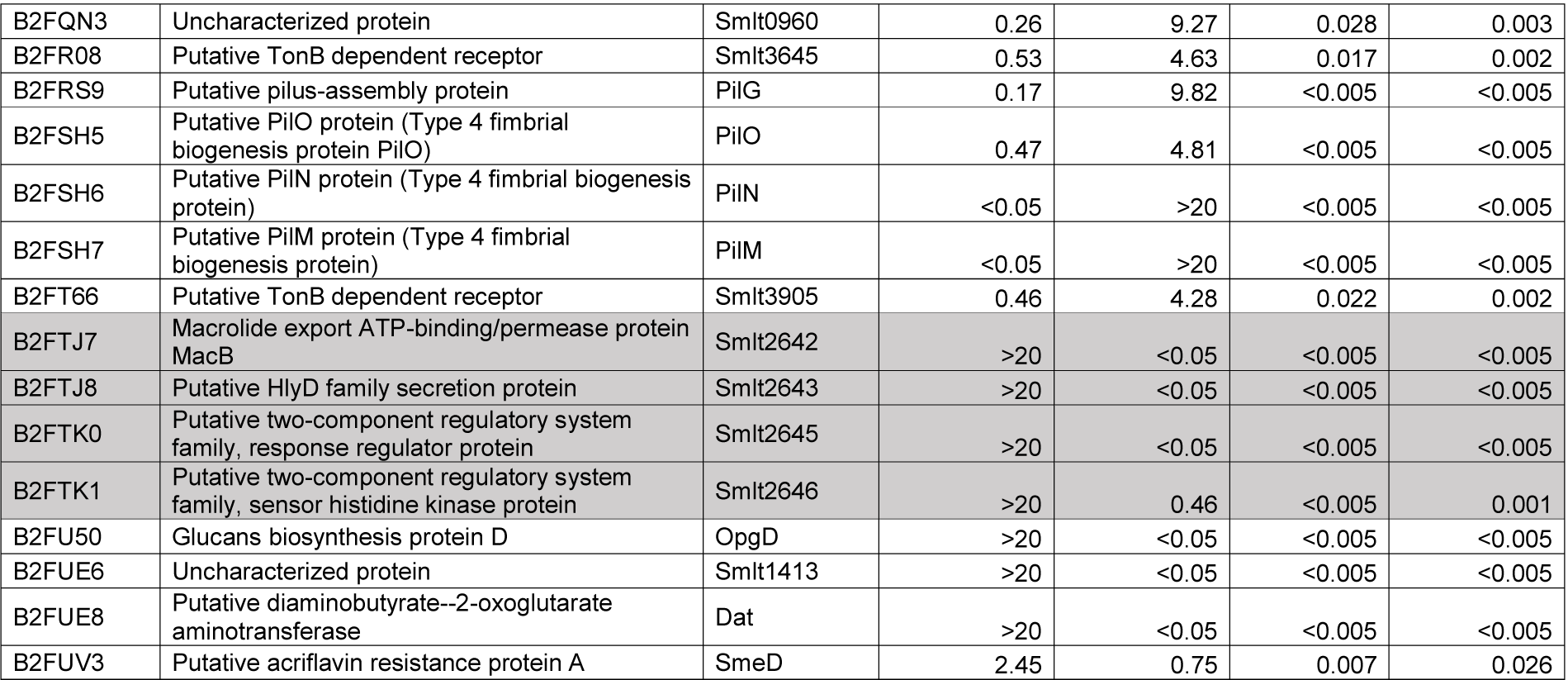
Significant changes in envelope protein abundance seen in *S. maltophilia* mutant K M6 LEV^R^ compared with K M6, which reverse upon disruption of sensor kinase gene *smlt2646*. Strains were grown in NB and fold changes in raw abundance are provided, averaged across three biological replicates of parent (K M6) and mutant (K M6 LEV^R^) and against parent (K M6 LEV^R^) and mutant (K M6 LEV^R^ *smlt2646*). Analysis was as described in Experimental and proteins listed are those with significantly up- or down-regulated abundance, (p <0.05) in K M6 LEV^R^ versus K M6, whose abundance was then significantly shifted back in the opposite direction in K M6 LEV^R^ *smlt2646* versus K M6 LEV^R^. Shaded proteins are those discussed in the text.

**Figure 2.**
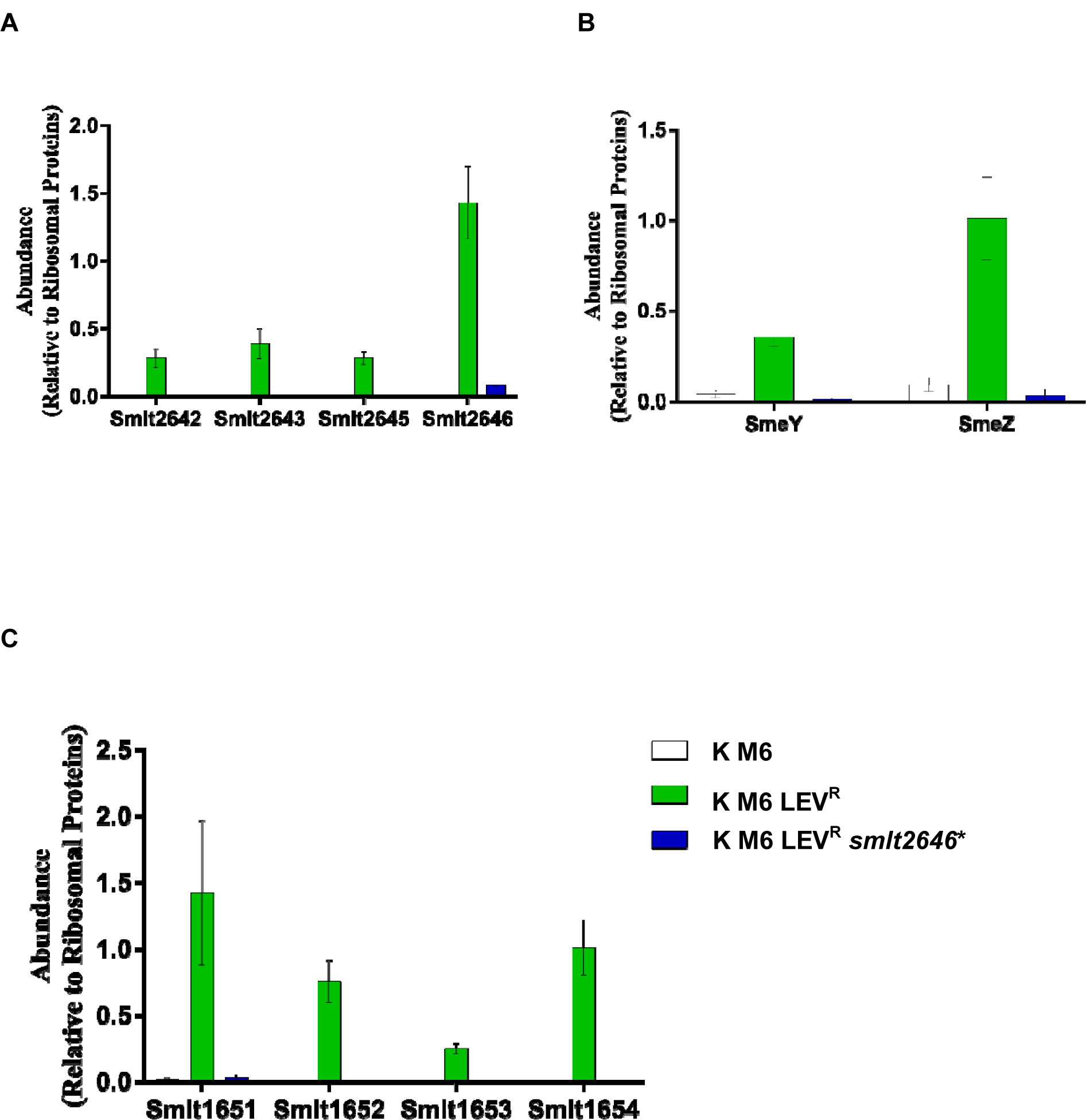
Impact of Smlt2646 sensor kinase activation on SmeYZ efflux production, and on Smlt2642/3 and Smlt1651-4 ABC transporter production. Protein abundance was measured using LC-MS/MS and normalised to the abundance of ribosomal proteins in cell extracts obtained from bacteria grown in NB. Data are mean ± standard error of the mean, *n*=3. Protein abundance in the mutant K M6 LEV^R^ is statistically significantly different from the parent strain and from the mutant where *smlt2646*\* was disrupted according to t-test (*p*<0.05). (**A**) Smlt2642/3 ABC transporter and Smlt2645/6 response regulator/sensor kinase production in the *smlt2546*\* activator mutant K M6 LEV^R^, and the *smlt2646*\* disrupted derivative versus parent strain K M6 (**B**) SmeYZ efflux pump production in the *smlt2546*\* activator mutant K M6 LEV^R^, and the *smlt2646*\* disrupted derivative versus parent strain K M6 (**C**) Smlt1651-4 ABC transporter production in the *smlt2546*\* activator mutant K M6 LEV^R^, and the *smlt2646*\* disrupted derivative versus parent strain K M6.

Since an activatory mutation in a two-component system is generally dominant *in trans*, we aimed to confirm the effect of the mutated version of the sensor kinase gene *smlt2646*, referred to as *smlt2646*\*, from K M6 LEV^R^ in a wild-type background. The operon, including the response regulator gene and the putatively activated sensor kinase mutant gene (*smlt2644-smlt2646*\*) from K M6 LEV^R^, was cloned to create plasmid pBBR1MCS-4::*smlt2644-6*\*, which was used to transform *S. maltophilia* K279aAmp^FS^, an ampicillin susceptible derivative of K279a (27) to ampicillin resistance (the marker on the plasmid). Relative to plasmid only control, MIC testing showed that carriage of pBBR1MCS-4::*smlt2644-6\** in K279aAmp^FS^ confers levofloxacin intermediate resistance, and a greatly increased MIC of amikacin (**Table 2**).

Disruption of the activated sensor kinase mutant gene *smlt2646*\* in K M6 LEV^R^ reduced Smlt2642/3 ABC transporter production back to the levels seen in K M6 (**Table 3, Figure 2A**) and reduced MICs of amikacin and levofloxacin to one doubling dilution below even their MICs against K M6 (**Table 2**). This confirms that the activator mutation seen in the sensor kinase Smlt2646* causes Smlt2642/3 ABC transporter upregulation and, together with the transactivation experiment, that the Smlt2646* mutation causes the resistance phenotype expressed by K M6 LEV^R^. However, disruption of the upregulated putative ABC transporter gene *smlt2642* in K M6 LEV^R^ only reduced the MIC of amikacin, and even then it remained two doubling dilutions higher than the MIC against K M6 (**Table 2**) showing that Smlt2642/3 transporter upregulation is not responsible for levofloxacin resistance in K M6 LEV^R^ and is only partially responsible for the increased MIC of amikacin against this mutant. In order to find additional amikacin resistance proteins, we explored the proteomics data (**Table 3**) and identified that the aminoglycoside efflux pump SmeYZ was also over-produced in K M6 LEV^R^ relative to K M6, then down regulated in upon disruption of the *smlt2646*\* sensor kinase gene in K M6 LEV^R^, i.e. its production mirrored changes in the MIC of amikacin (**Figure 2B, Table 2**). Therefore, we conclude that increased amikacin MIC seen when the Smlt2645/6 two-component system is activated by mutation is caused by a combined effect of SmeYZ and Smlt2642/3 over-production. However, neither Smlt2642/3 (**Table 2**) or SmeYZ (14) are responsible for levofloxacin resistance in K M6 LEV^R^ so we again searched the proteomics data (**Table 3**) and identified another novel ABC transporter, Smlt1651-4, which was upregulated in K M6 LEV^R^ relative to K M6 and then downregulated in the *smlt2646*\* signal sensor gene disrupted derivative of K M6 LEV^R^ (**Figure 2C**), i.e. a derivative that lost levofloxacin resistance (**Table 2**). We therefore disrupted the putative ABC transporter gene *smlt1651* in K M6 LEV^R^ and noted that the MIC of levofloxacin reduced to be the same as the MIC against K M6, but the amikacin MIC did not change (**Table 2**). This confirmed that over-production of Smlt1651-4 is responsible for levofloxacin resistance in K M6 LEV^R^. SmeDEF over-production, seen in K M6 and maintained in K M6 LEV^R^ (**Figure 1A**) is also essential for levofloxacin resistance in K M6 LEV^R^ as confirmed because disruption of *smeE* reduced the levofloxacin MIC against K M6 LEV^R^ even more than disruption of the ABC transporter gene *smlt1651* (**Table 2**). Importantly, however, the MIC of levofloxacin against K M6 LEV^R^ *smeE* remained one doubling dilution higher than against K M6 *smeE* (**Table 2**) confirming involvement of ABC transporter Smlt1651-4 over-production in elevating levofloxacin MICs in *S. maltophilia*.

### Conclusions

Over-production of SmeDEF confers levofloxacin resistance in *S. maltophilia* (22). This is typically caused by an *smeT* loss-of-function mutation, as seen here in K279a derived mutant K M7 (**Table 2**). However, we have also found a novel alternative mutational pathway to this phenotype. We show that disruption of the glycosyl transferase gene *smlt0622* constitutively activates production of SmeDEF (**Figure 1B**). A loss-of-function mutation in this gene has a significant impact of cell growth (**Figure 1D**), but the laboratory selected *smlt0622* point mutant, K M6 appears to retain some residual Smlt0622 activity, because SmeDEF production is not at such high levels (**Figure 1B**) and growth rate is not significantly affected (**Figure 1D**). We hypothesise that reduction of Smlt0622 activity affects the concentration of some cellular metabolite, possibly increasing the concentration of a toxic molecule that is a signal for SmeT activation. This would imply there are multiple signals for SmeT de-repression since it is known that triclosan can also perform this role (18). It may be that, like triclosan, the putative cytoplasmic SmeT-activator ligand is also a substrate for SmeDEF. In this way, the SmeT-SmeDEF regulatory system may be analogous to the VceCAB efflux pump and its control by the SmeT homologue VceR in *Vibrio cholerae*, where VceR can be de-repressed in the presence of a number of different substrates of VceCAB (28,29). Testing this hypothesis will form the basis of future work.

Because SmeDEF abundance is not increased to the same extent in the *smlt0622* point mutant K M6 as it is in the *smeT* loss-of-function mutant K M7 (**Figure 1A**) the MIC of levofloxacin against K M6 is not high enough for the mutant to be called resistant (**Table 2**). Therefore, by selecting a resistant derivative, K M6 LEV^R^, we were able to identify a novel two-component regulatory system Smlt2645/6, where Smlt2646 is a sensor histidine kinase and Smlt2645 is a response regulator. Activation of the Smlt2646 sensor kinase by mutation increases production of two novel ABC-type antibiotic efflux pumps, and the known aminoglycoside efflux pump SmeYZ (14). Alongside SmeYZ over-production, amikacin MICs increased in K M6 LEV^R^ because of the over-production of the novel ABC transporter Smlt2642/3 (**Figure 2**) as annotated in the *S. maltophilia* K279a genome sequence (30). We now name this novel *<u>S</u>. <u>m</u>altophilia* <u>A</u>BC transporter: “SmaAB”. The Smlt2645/6 two-component system encoded immediately adjacent to *smaAB*, we name SmaRS. A second novel ABC transporter, Smlt1651-4, which we now name SmaCDEF, is also up-regulated upon activation of the SmaRS two-component system (**Figure 2**), and this enhances the MIC of levofloxacin (but not amikacin), and when this occurs in addition to SmeDEF over-production, this confers levofloxacin resistance (**Table 2**).

Accordingly, we have added to the already dizzying array of known efflux systems relevant for intrinsic and acquired antimicrobial resistance in *S. maltophilia* (31). A species having a remarkable resistance protein armamentarium, explaining why it is one of the most difficult-to-treat bacterial pathogens.

## Experimental

### Materials, bacterial isolates and antimicrobial susceptibility testing

Chemicals were from Sigma and growth media from Oxoid, unless otherwise stated. Strains used were *S. maltophilia* K279a (32) two spontaneous mutants selected for reduced moxifloxacin susceptibility, K M6 and K M7 (19) and a β-lactam susceptible mutant derivative, K279a *ampR*^FS^ with a frameshift mutation engineered into the β-lactamase activator gene *ampR* via suicide gene replacement (27). Antimicrobial susceptibility was determined using CLSI broth microtiter assays (33) or disc susceptibility testing (34) and interpreted using published breakpoints (1).

### Selection and construction of mutants

To select levofloxacin resistant mutant derivative of K M6, 100 µL aliquots of overnight cultures of K M6 grown in Nutrient Broth (NB) were spread onto Mueller Hinton agar containing 5 µg.mL^-1^ levofloxacin and incubated for 24 h. Insertional inactivation of *smlt0622, smlt2646*\*, *smlt2643, smlt1651* and *smeE* was performed using the pKNOCK suicide plasmid (35). The DNA fragments were amplified with Phusion High-Fidelity DNA Polymerase (NEB, UK) from *S. maltophilia* K279a genomic DNA. pKNOCK-GM::*smeE* was constructed by PCR using primers *smeE* F (5′-CAATGTTGTCGATCGCCTGA-3′) and *smeE* R (5′-TACGACATCGCCGTCCATTC-3′), the product was digested with PstI and XhoI and ligated into pKNOCK-GM at the PstI and XhoI sites. pKNOCK-GM::*smlt0622* was constructed by using *smlt0622* F (5′-CAACGAGCGGGATGTTAGGT-3′) and *smlt0622* R (5′-CGTCGAAGTGGGCAACAAC-3′), the product was digested with BamHI and XhoI and ligated into pKNOCK-GM at the BamHI and XhoI sites. pKNOCK-GM::*smlt1651*, pKNOCK-GM::*smlt2643* and pKNOCK-GM::*smlt2646* were constructed using primers *smlt1651* FW KO with a SalI site included, underlined (5′-AAA<u>GTCGACA</u>GTGGTGGAAGGTGCTGG-3′) and *smlt1651* RV KO with ApaI (5′-AAA<u>GGGCCC</u>GGCATGGAAGTAGGTATCGACA-3′); *smlt2643* FW KO with SalI (5′-AAA<u>AGTCGA</u>CCCACAGTGGCTCCAAGAAAC-3′) and *smlt2643* RV KO with ApaI (5′-ATA<u>GGGCCC</u>GGCATCATCACTTTCGGCAA-3′); *smlt2646* FW KO with SalI (5′-AAA<u>GTCGAC</u>TATGACGAGCCGGAAACCAT-3′) and *smlt2646* RV KO with ApaI (5′-AAA<u>GGGCCC</u>CCATGGAGTTGAAGTCGCTG-3′). Each recombinant plasmid was then transferred into K279a, K M6 or K M6 LEV^R^, as required, by conjugation from *Escherichia coli* BW20767. Mutants were selected using gentamicin (30 µg.mL^-1^) and the mutations were confirmed by PCR using primers *smeE* F and *smeE* R (above); *smlt0622* F and *smlt0622* R (above); *smlt1651* F (5′-AGAGCAGGTGGGGGCGTCTGAACGCC-3′) and BT543 (5′-TGACGCGTCCTCGGTAC-3′); *smlt2643* F (5′-CTGCAGGCATGAGACTCAGT-3′) and BT543; *smlt2646* F (5′-TTGCAGGACCGGGTGGACGCAACG-3′) and BT543.

### Proteomics

500 µL of an overnight NB culture were transferred to 50 mL NB and cells were grown at 37°C to 0.6 OD_600_. Cells were pelleted by centrifugation (10 min, 4,000 × *g*, 4°C) and resuspended in 30 mL of 30 mM Tris-HCl, pH 8 and broken by sonication using a cycle of 1 s on, 0.5 s off for 3 min at amplitude of 63% using a Sonics Vibracell VC-505TM (Sonics and Materials Inc., Newton, Connecticut, USA). The sonicated samples were centrifuged at 8,000 rpm (Sorval RC5B PLUS using an SS-34 rotor) for 15 min at 4°C to pellet intact cells and large cell debris; For envelope preparations, the supernatant was subjected to centrifugation at 20,000 rpm for 60 min at 4°C using the above rotor to pellet total envelopes. To isolate total envelope proteins, this total envelope pellet was solubilised using 200 μL of 30 mM Tris-HCl pH 8 containing 0.5% (w/v) SDS.

Protein concentrations in all samples were quantified using Biorad Protein Assay Dye Reagent Concentrate according to the manufacturer’s instructions. Proteins (5 µg/lane for envelope protein analysis) were separated by SDS-PAGE using 11% acrylamide, 0.5% bis-acrylamide (Biorad) gels and a Biorad Min-Protein Tetracell chamber model 3000×1. Gels were resolved at 200 V until the dye front had moved approximately 1 cm into the separating gel. Proteins in all gels were stained with Instant Blue (Expedeon) for 20 min and de-stained in water.

The 1 cm of gel lane was subjected to in-gel tryptic digestion using a DigestPro automated digestion unit (Intavis Ltd). The resulting peptides from each gel fragment were fractionated separately using an Ultimate 3000 nanoHPLC system in line with an LTQ-Orbitrap Velos mass spectrometer (Thermo Scientific). In brief, peptides in 1% (v/v) formic acid were injected onto an Acclaim PepMap C18 nano-trap column (Thermo Scientific). After washing with 0.5% (v/v) acetonitrile plus 0.1% (v/v) formic acid, peptides were resolved on a 250 mm × 75 μm Acclaim PepMap C18 reverse phase analytical column (Thermo Scientific) over a 150 min organic gradient, using 7 gradient segments (1-6% solvent B over 1 min, 6-15% B over 58 min, 15-32% B over 58 min, 32-40% B over 5 min, 40-90% B over 1 min, held at 90% B for 6 min and then reduced to 1% B over 1 min) with a flow rate of 300 nL/min. Solvent A was 0.1% formic acid and Solvent B was aqueous 80% acetonitrile in 0.1% formic acid. Peptides were ionized by nano-electrospray ionization MS at 2.1 kV using a stainless-steel emitter with an internal diameter of 30 μm (Thermo Scientific) and a capillary temperature of 250°C. Tandem mass spectra were acquired using an LTQ-Orbitrap Velos mass spectrometer controlled by Xcalibur 2.1 software (Thermo Scientific) and operated in data-dependent acquisition mode. The Orbitrap was set to analyse the survey scans at 60,000 resolution (at m/z 400) in the mass range m/z 300 to 2000 and the top twenty multiply charged ions in each duty cycle selected for MS/MS in the LTQ linear ion trap. Charge state filtering, where unassigned precursor ions were not selected for fragmentation, and dynamic exclusion (repeat count, 1; repeat duration, 30 s; exclusion list size, 500) were used. Fragmentation conditions in the LTQ were as follows: normalized collision energy, 40%; activation q, 0.25; activation time 10 ms; and minimum ion selection intensity, 500 counts.

The raw data files were processed and quantified using Proteome Discoverer software v1.4 (Thermo Scientific) and searched against the UniProt *S. maltophilia* strain K279a database (4365 protein entries; UniProt accession UP000008840) using the SEQUEST (Ver. 28 Rev. 13) algorithm. Peptide precursor mass tolerance was set at 10 ppm, and MS/MS tolerance was set at 0.8 Da. Search criteria included carbamidomethylation of cysteine (+57.0214) as a fixed modification and oxidation of methionine (+15.9949) as a variable modification. Searches were performed with full tryptic digestion and a maximum of 1 missed cleavage was allowed. The reverse database search option was enabled, and all peptide data was filtered to satisfy false discovery rate (FDR) of 5 %. Protein abundance measurements were calculated from peptide peak areas using the Top 3 method (36) and proteins with fewer than three peptides identified were excluded. The proteomic analysis was repeated three times for each parent and mutant strain, each using a separate batch of cells. Data analysis was as follows: all raw protein abundance data were uploaded into Microsoft Excel. Raw data from each sample were normalised by division by the average abundance of all 30S and 50S ribosomal protein in that sample. A one-tailed, unpaired T-test was used to calculate the significance of any difference in normalised protein abundance data in the three sets of data from the parent strains versus the three sets of data from the mutant derivative. A *p*-value of <0.05 was considered significant. The fold change in abundance for each protein in the mutant compared to its parent was calculated using the averages of normalised protein abundance data for the three biological replicates for each strain.

### Whole genome sequencing to Identify mutations

Whole genome resequencing was performed by MicrobesNG (Birmingham, UK) on a HiSeq 2500 instrument (Illumina, San Diego, CA, USA). Reads were trimmed using Trimmomatic (37) and assembled into contigs using SPAdes 3.10.1 (http://cab.spbu.ru/software/spades/). Assembled contigs were mapped to *S. maltophilia* K279a (30) obtained from GenBank (accession number NC_010943) by using progressive Mauve alignment software (38).

### Cloning smlt2644-6 for in trans expression

*In trans* expression of Smlt2646* was performed after amplifying the *smlt2644-6* operon with Phusion High-Fidelity DNA Polymerase (NEB, UK) using K M6 LEV^R^ genomic DNA and primers *smlt2644* F with an EcoRI site added, underlined, (5′-AAA<u>GAATTC</u>TTGGAGCCACTGTGGAGATTG-3′) and *smlt2646* R with EcoRI (5′-AAA<u>GAATTC</u>GGTGGGTCGGGGGTAGAGT-3′). The resulting DNA was digested with EcoRI and ligated to pBBR1MCS-4 at its EcoRI site (39,40). Recombinant plasmid was then transferred into K279a *ampR*^FS^ by electroporation. K279a *ampR*^FS^/pBBR1MCS-4 and K279a *ampR*^FS^/pBBR1MCS-4::*smlt2644-6* were selected using ampicillin (100 µg.mL^-1^) and the presence of plasmids were confirmed by PCR using primers M13F (5′-GTAAAACGACGGCCAGT-3′) and M13R (5′-CAGGAAACAGCTATGAC-3′).

### Growth curves

OD_600_ measurements of bacterial cultures were performed using a Spectrostar Nano Microplate Reader (BMG, Germany) in COSTAR Flat Bottom 96-well plates. Overnight cultures (in NB) were adjusted to OD_600_ = 0.01 and 200 µL of the diluted culture were taken to the plate together with a blank, NB. The plate was incubated at 37°C with double orbital shaking and OD_600_ was measured every 10 min for 24 h.

## Acknowledgments

This work was funded by grant MR/S004769/1 to M.B.A. from the Antimicrobial Resistance Cross Council Initiative supported by the seven United Kingdom research councils and the National Institute for Health Research. K.C. received a postgraduate scholarship from SENESCYT, Ecuador. Genome sequencing was provided by MicrobesNG (http://www.microbesng.uk/), which is supported by the BBSRC (grant number BB/L024209/1).

**We declare no conflicts of interest**.

